# Patient-derived endometrial organoids from MRKH patients: Insight in disease causing pathways

**DOI:** 10.1101/2021.10.27.466065

**Authors:** Sara Y. Brucker, Thomas Hentrich, Julia M. Schulze-Hentrich, Martin Pietzsch, Noel Wajngarten, Anjali Ralhan Singh, Katharina Rall, André Koch

**Author notes:** These authors contributed equally (Sara Y. Brucker, Thomas Hentrich, Julia M. Schulze-Hentrich).

## Abstract

The uterus is responsible for the nourishment and mechanical protection of the developing embryo and fetus and is an essential part in mammalian reproduction. The Mayer-Rokitansky-Küster-Hauser (MRKH) syndrome is characterized by agenesis of the uterus and upper part of the vagina in females with normal ovarian function. Although heavily studied, the cause of the disease is still enigmatic. Current research in the field of MRKH mainly focusses on DNA-sequencing efforts and, so far, failed to decipher the nature and heterogeneity of the disease, thereby holding back scientific and clinical progress. Here, we developed long-term expandable organoid cultures from endometrium found in uterine rudiment horns of MRKH patients. Phenotypically, they share great similarity with healthy control organoids and are surprisingly fully hormone responsive. Transcriptome analyses, however, identified an array of dysregulated genes that point at potentially disease-causing pathways altered during the development of the female reproductive tract. We consider the endometrial organoid cultures to be a powerful research tool that promise to enable an array of studies into the pathogenic origins of MRKH syndrome and possible treatment opportunities to improve patient quality of life.

## Introduction

The Mayer-Rokitansky-Küster-Hauser (MRKH) syndrome (OMIM 277000) is a rare malformation, characterized by the partial or complete absence of the uterus and the upper two-thirds of the vagina, due to a still unknown defect in embryonic development (Oppelt et al., 2006). It affects 1 in 4,500 women, making it the second most common reason for primary amenorrhea (Aittomaki et al., 2001, Herlin et al., 2016, Timmreck and Reindollar, 2003). Typically, MRKH patients have a normal female karyotype (46, XX) and regular development of secondary sexual characteristics as their ovaries are functional. While the MRKH syndrome can occur as an isolated genital malformation (type 1), it is often associated with additional renal and/or skeletal, and to a lesser extent with auditory, cardiac and other defects (type 2) (Oppelt et al., 2012, Rall et al., 2015b). In both cases, patients with MRKH frequently have one or two uterine remnants, consisting of myometrium and less often also endometrium (Rall et al., 2013).

The etiology of the syndrome remains largely enigmatic, yet the spectrum of malformations encountered in MRKH patients suggests the disease to originate from a developmental defect during embryogenesis. Moreover, cases of familial clustering have implied a genetic component in the etiology (Herlin et al., 2014, Williams et al., 2017, Nik-Zainal et al., 2011). Mutations in *WNT4*, which is essential for the complete formation of the Müllerian ducts (Vainio et al., 1999), have been reported earlier as a possible cause for MRKH in a small number of patients (Biason-Lauber et al., 2004, Philibert et al., 2008, Philibert et al., 2011). Since WNT4 is also necessary to prevent formation of Leydig cells in women, patients with mutated WNT4 also present with clinical hyperandrogenism, rendering it a slightly different clinical entity (Biason-Lauber et al., 2007, Biason-Lauber et al., 2004, Philibert et al., 2008). Furthermore, in some patients, mutations and possibly harmful variants have been found in developmental genes like *WNT9B* (Wang et al., 2014, Waschk et al., 2016), *LHX1* (Ledig et al., 2012, Ledig et al., 2011, Sandbacka et al., 2013), or *TBX6* (Sandbacka et al., 2013, Tewes et al., 2019, Tewes et al., 2015). Of high interest are specifically *LHX1, WNT4*, and *WNT9B* due to their previously reported role in formation of the Müllerian Ducts (MD) from the coelomic epithelium in gestational week six (Mullen and Behringer, 2014). After the two MDs are formed, they start growing caudally along the Wolffian Ducts. By week eight, the terminal ends of the MDs begin to fuse, forming the uterovaginal duct. In males, the MDs start to regress after week ten under the influence of AMH and WNT7A. In females, however, they differentiate into fallopian tubes, uterus, cervix, and vagina specifically regulated by the coordinated action of transcription factors and signaling molecules such as homeodomain transcription factors (e.g., Hox genes) and members of the WNT family (Robboy et al., 2017, Roly et al., 2018). This concerted interplay between transcription factors, hormones, and growth signals during embryogenesis leads to a variety of highly timed and spatial gene expression changes (Roly et al., 2020). The widespread lack of a clear genetic link in MRKH syndrome suggests that the answer might be found at the transcriptional level. Therefore, the focus in recent years has shifted towards molecular characterizations of primary diseased tissue, identifying a plethora of new candidate genes and regulatory networks that might drive or contribute to the pathology (Hentrich et al., 2020, Rall et al., 2015a). In addition, hormonal stimulation of primary endometrial stroma cultures derived from MRKH rudimentary tissue showed a reduced transcriptional response compared to healthy controls(Brucker et al., 2017), indicating that dysfunctional hormone receptors play a role in the pathophysiology of MRKH (Rall et al., 2013). However, as other estrogen and progesterone dependent tissues like the breast are normally developed in MRKH patients, a possible defect would have to be limited to the genital tract. This stresses the importance of studying the rudimentary tissue directly. Whereas stromal cultures have already been investigated (Brucker et al., 2017), the absence of a functional model for the glandular epithelium of MRKH endometrium to analyze the characteristics and capabilities of affected cell types vastly limited the pathophysiological understanding of this disease. Recent years have seen the establishment of patient-derived organoid models from healthy and diseased endometrium (Turco et al., 2017, Boretto et al., 2017, Boretto et al., 2019). Organoids are self-renewing, three dimensional (3D) models that mimic key properties and characteristics of the original *in vivo* tissue which greatly facilitates research into complex interactions of involved cell types.

Here, we show for the first time the successful establishment of organoid models from endometrium found in rudimentary tissue of MRKH patients. The established organoids showed high similarity to healthy endometrial organoids, were hormone-responsive, and could be cultured for long-term. Yet, gene expression analyses by RNA-sequencing showed distinctive differences between diseased and healthy organoids, emphasizing their potential as a powerful tool to investigate the etiology of the MRKH syndrome. We identified several important developmental transcription factors as differentially expressed between healthy and MRKH organoids. This highlights the importance and value of these models for studying and understanding of the pathogenesis. The establishment and long-term growth of epithelial tissue from MRKH rudiments provides a powerful addition to the toolbox for studying the disease in a controlled and standardized environment, and these endometrial organoids promise to pave new avenues towards a better understanding and possible treatments of the disease.

## Results

### Long-term 3D organoid cultures can be established from uterine rudiment endometrium

Human endometrial organoid cultures have recently been established by several research groups (Turco et al., 2017, Boretto et al., 2019, Boretto et al., 2017). Typically, glandular structures obtained after processing endometrial biopsies are embedded in an extracellular matrix component such as BME or Matrigel and cultured in presence of a defined cocktail of growth factors (Supplementary Table 2). Since endometrium can be found in MRKH uterine rudiments (Rall et al., 2013), we screened a cohort of MRKH patients that underwent a laparoscopically assisted creation of a neovagina for the presence of endometrium in uterine rudiments. A total of 48 patients (35 MRKH-Type I (73%), 13 MRKH-Type II (27%)) were screened of which 37 (32/35 Type I (91%), 5/13 Type II (38%)) had uterine rudiments present (Fig. 1a). We were able to macroscopically detect endometrium (Fig. 1b, c) in 12 patients (12/32 Type I (33%), 0/5 Type II (0%)). Endometrium of four rudiments was also subjected to histological processing (Fig. 1b, Asterisk) and immunohistochemistry for the transcription factor Pax8 (highly expressed in endometrial epithelial cells), estrogen receptor alpha (ER), progesterone receptor (PR) and the proliferation marker Ki67/Mib1 (Fig. 1d), to verify the endometrium entity of the tissue. Pax8 staining confirmed that endometrial epithelium of MRKH patients had all morphological features of normal endometrium showing tubular and frequently branching glands with a single layer of columnar epithelium (Fig. 1d) as it has been shown recently in an extensive histology study of MRKH rudiments (Rall et al., 2013). Whereas ER and PR were ubiquitously expressed in the epithelial as well as stromal cells of the endometrium, the proliferative capacity, measured by Ki67/Mib1 expression, was almost absent in epithelium and stroma of MRKH patients (Fig. 1d), as reported before (Rall et al., 2013). Although the proliferative capacity of the initial cell population was low, we successfully established 14 organoid models (ten patients had endometrium present unilaterally and two patients bilaterally, which accounts for the two additional organoid models) (Supplementary Table 1). The success rate in establishing organoid models from patients with macroscopically detected endometrium was 100% (14/14). The fact that the endometrium size of MRKH patients ranged from a few millimeters in diameter (e.g., MRKH #03 – Fig. 1d) to even under a millimeter (e.g., MRKH #04 – Fig. 1d) limited the amount of starting material for culture setup but did not hinder the successful establishment of an organoid model (compare MRKH #03 and #04, Fig. 1e). Within 3 – 6 days, cystic organoid structures were visible, and all 14 established MRKH organoid models as well as four control models (Fig. 1e, Supplementary Fig. 1), were successfully cultured long-term for more than 15 passages (>6 months of culture) without showing signs of a decrease in proliferation.

**Figure 1.**
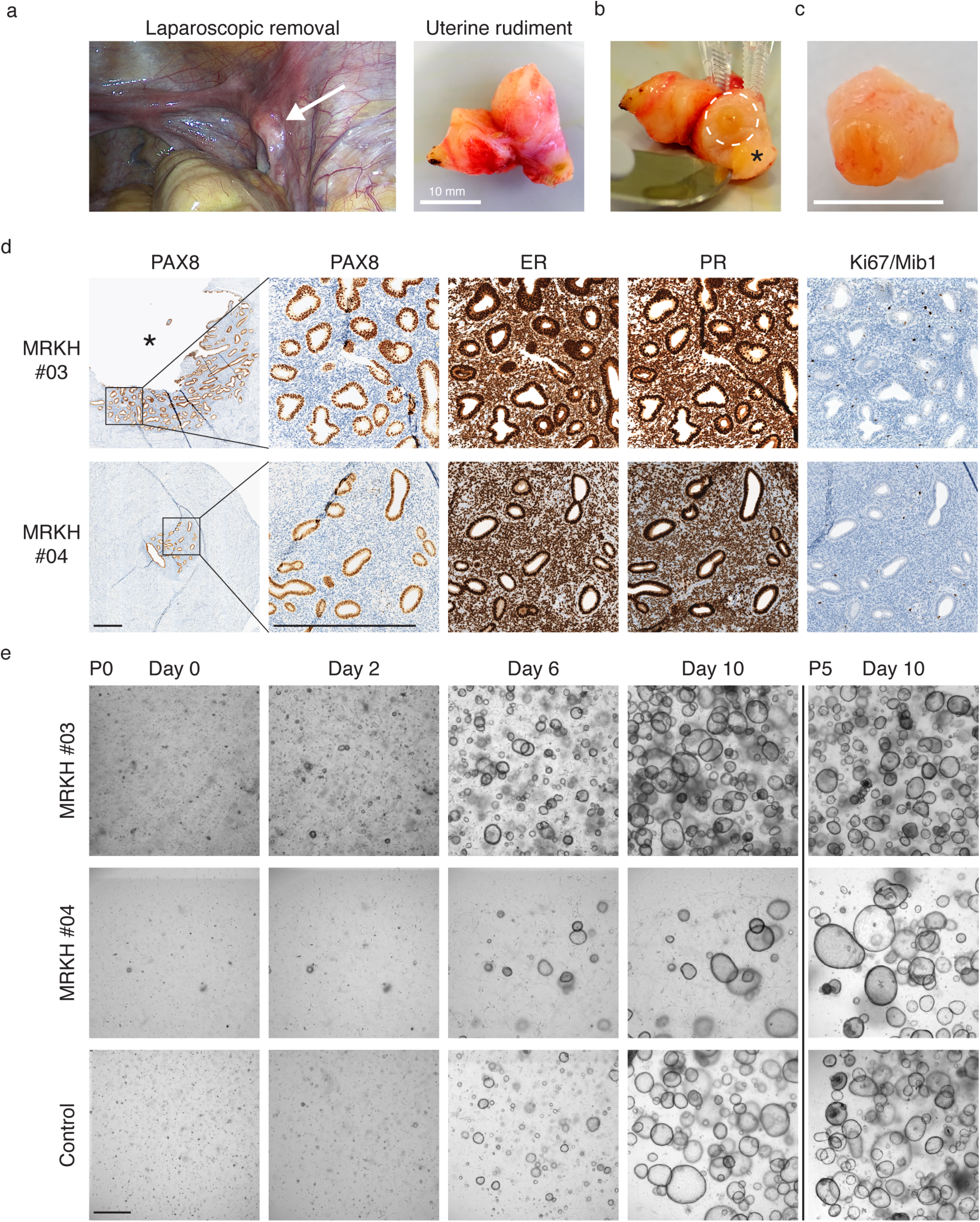
Long-term 3D organoid cultures can be established from uterine rudiment endometrium. **a)** Left: Laparoscopic image of an 18-year-old patient with MRKH syndrome (MRKH #03). White arrow indicates the uterine rudiment horn. Right: Laparoscopically excised uterine rudiment horn before segmentation; Scale bar = 10 mm **b)** Sectioned uterine rudiment horn of MRKH #03. White circle depicts macroscopical endometrium used for organoid establishment (see c). Asterisk indicates part that underwent pathological characterization (see corresponding pictures in d) **c)** Excised section of macroscopical endometrium used for digestion and organoid establishment; Scale bar = 10 mm **d)** Immunohistochemical characterization of uterine rudiment from MRKH-patient #03 (top row) and MRKH-patient #04 (bottom row). Analyzed were PAX8, Estrogen receptor alpha (ER), Progesterone receptor (PR), and proliferation marker Ki67/Mib1. The asterisk in MRKH #03 indicates the region marked in b). Both models show endometrial gland structures (PAX8-positive) with widespread and intense ER and PR expression in glandular and stromal compartments. There is almost no proliferation capacity visible in both MRKH tissues; Scale bar = 500 μm **e)** Bright-field images of cell suspensions from endometrial MRKH tissue digestions (Top and middle row) as well as from a healthy control (bottom) after seeding (P0). Organoid growth for the same spot on the culture plate was monitored over the course of ten days (Day 0-10). The right panel shows the same cultures at day 10 of the fifth passage (P5). Scale bar = 500 μm

### MRKH organoids show high phenotypic similarity to organoids from healthy controls

The endometrial organoids of MRKH patients and healthy controls were expanded and passaged at ratios of 1:2 or 1:3 every 10 – 14 days and characterized by immunohistochemistry and immunofluorescence (Fig. 2a). Similar to the primary tissue, the endometrial organoids showed high and ubiquitous expression of Pax8 and ER, whereas the absence of steroid hormones (e.g., estradiol) in culture medium concludes to the absence of PR expression in all models. In sharp contrast to the observations in MRKH endometrial tissue, organoids of MRKH patients expressed high levels of the proliferation marker Ki67/Mib1, comparable to healthy controls. Markers of glandular epithelium (Cytokeratin, EpCAM, and E-Cadherin) were ubiquitously expressed in all organoid models and Perlecan staining at the basolateral membrane throughout the entirety of the endometrial organoids showed that epithelial polarity remained intact (Fig. 2b). Interestingly, phenotypic and morphological characterisations of healthy and diseased organoid models revealed no obvious differences.

**Figure 2.**
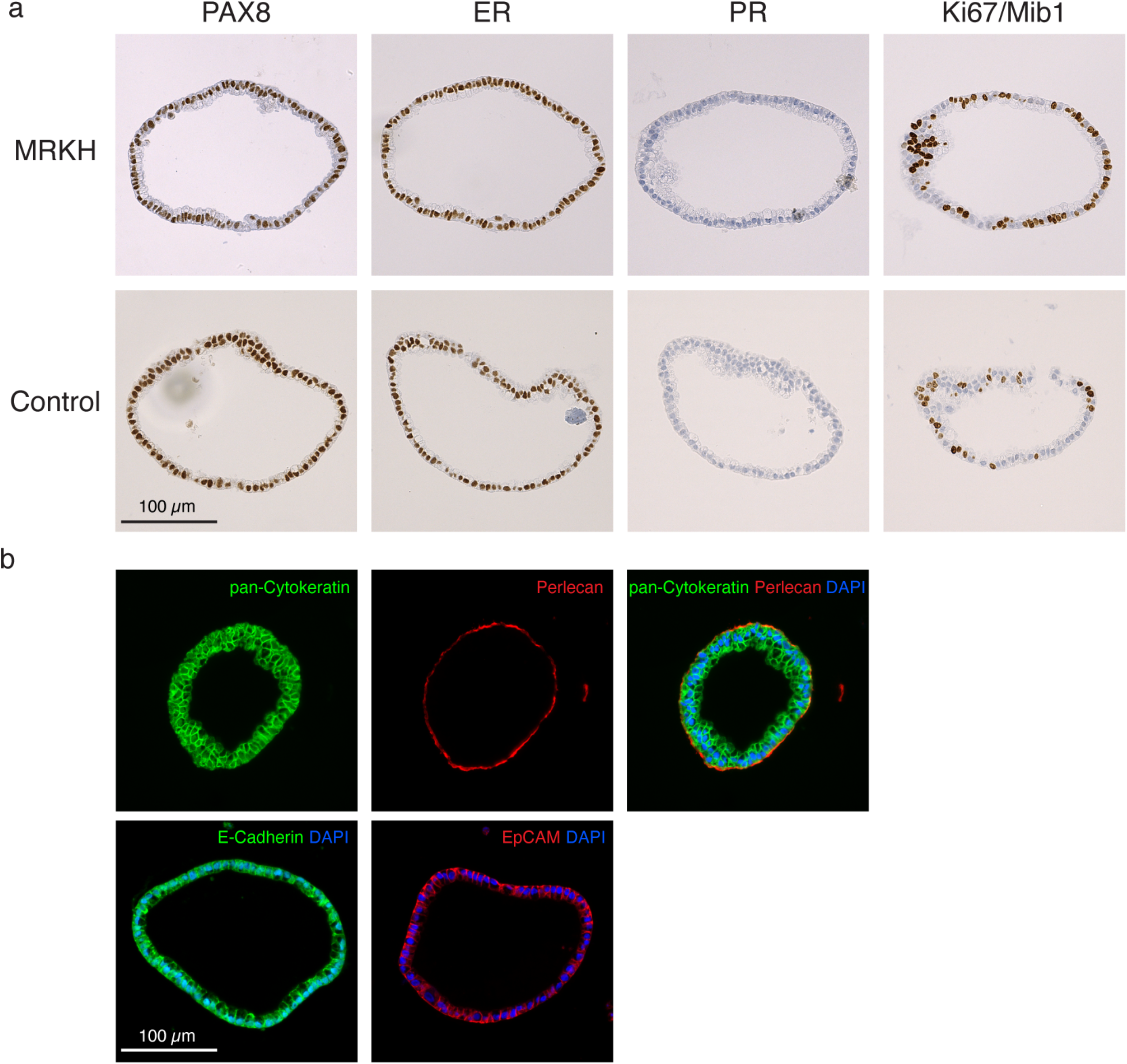
MRKH endometrial organoids show high phenotypic similarity to endometrial organoids from healthy controls. **a)** Representative immunohistochemistry images of sections from FFP-embedded MRKH (Top row) and control (Bottom row) endometrial organoids. Analyzed were the transcription factor PAX8, Estrogen receptor alpha (ER), Progesterone receptor (PR), and proliferation marker Ki67/Mib1. Scale bar = 100 μm **b)** Representative immunofluorescence images of sections from FFP-embedded MRKH endometrial organoids. Top row: Epithelial origin of organoids is shown by pan-Cytokeratin (green); epithelial polarity is shown by Perlecan (red), Nuclei stained with DAPI (blue). Bottom Row: Epithelial markers E-Cadherin (Left, green) and EpCAM (right, red) are shown in independent stainings of MRKH organoids. DAPI (blue) used as counterstain for nuclei. Scale bar = 100 μm

### Organoids derived from MRKH patients differ transcriptionally from healthy controls

After investigating organoid morphology, we next interrogated their transcriptome. Using RNA-sequencing, we profiled seven organoid models from MRKH patients and four from healthy controls that were grown in expansion medium (ExM), treated with beta-estradiol (E2) or the combination of beta-estradiol and progesterone (P4). Based on these experimental groups, differential gene expression was determined for the primary contrasts with cut-offs of *p*_BH_ ≤ 0.05 and |log_2_*FC*| ≥ 0.5 (Fig. 3a, Supplementary Table 5).

**Figure 3.**
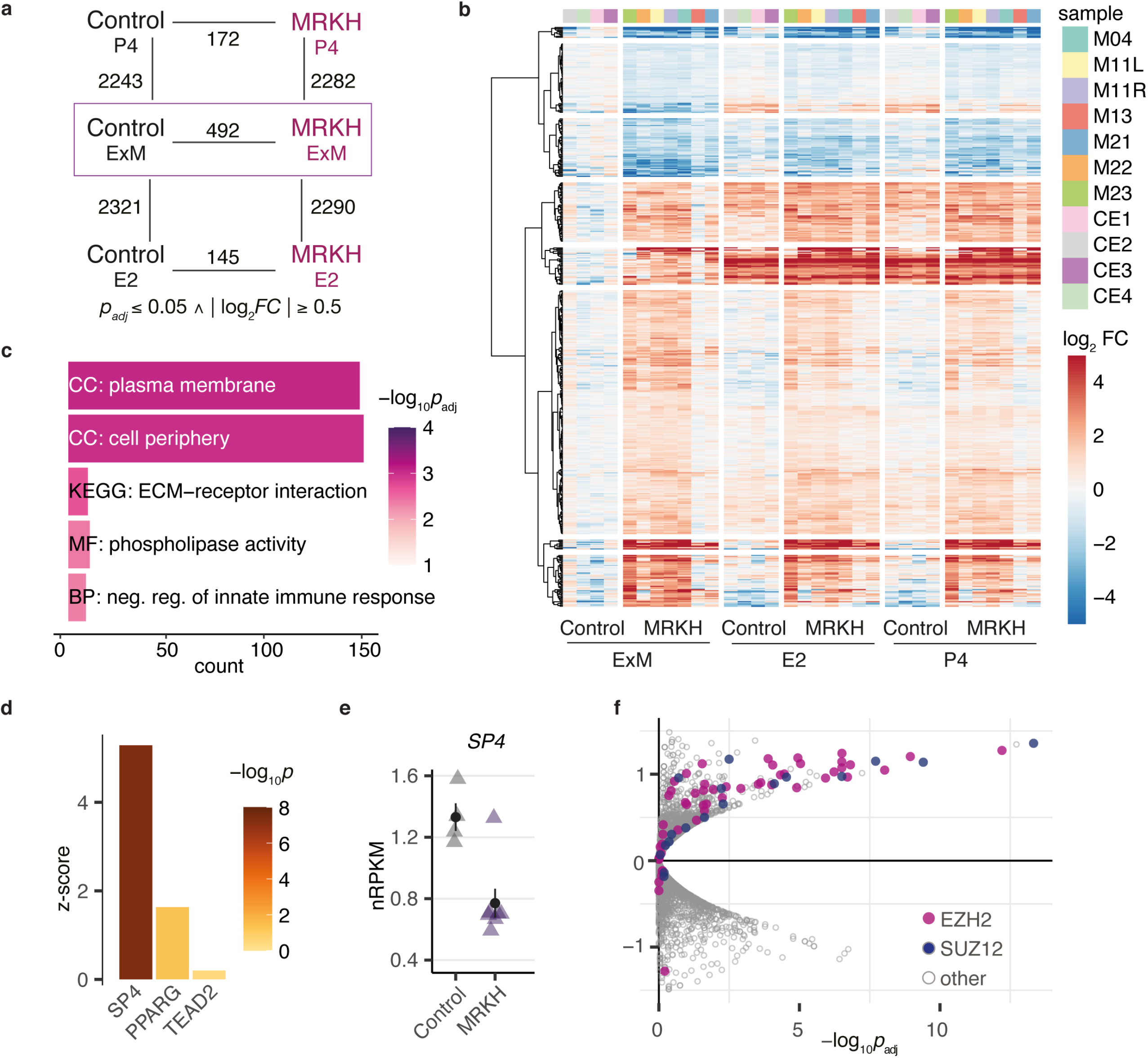
Organoids derived from MRKH patients and healthy controls differ transcriptionally. **a)** Diagram of experimental groups and number of differentially expressed genes (DEGs) between pairwise comparisons according to indicated fold-change and significance cut-offs. Control: organoids derived from unaffected women; MRKH: organoids derived from MRKH patients; ExM: organoids grown in expansion medium; E2: organoids treated with beta-estradiol; P4: organoids treated with beta-estradiol and progesterone. **b)** Expression profiles (*log_2_* expression change relative to Control ExM group) of 492 DEGs across all samples. Rows hierarchically clustered by Euclidian distance and ward.D2 method. Patient origin color-coded on top. **c)** Enrichment analysis of overrepresented Gene Ontology and KEGG terms for 492 DEGs (identified in MRKH/Control in ExM). Top five most significant terms with number of associated genes shown. CC: cellular compartment, MF: molecular function, BP: biological process. **d)** Transcription factor binding site analysis of 492 DEGs. Depicted are top three scoring position weight matrices of transcription factors that are also differentially expressed. Higher z-scores reflect higher enrichment of the binding motif among DEGs. **e)** Expression levels for SP4 plotted as individual data points with mean ± SEM. **f)** Enrichment analysis of transcriptional regulators for 492 DEGs identified in MRKH organoids based on ChIP-seq and DNase-I data according to TFEA.ChIP. EZH2 and SUZ12 are predicted to be significantly overrepresented among DEGs relative to the genome. Analysis based on default parameters for binding sites <1kB upstream including enhancer elements. Each dot represents a ChIP-seq accession, EZH2- and SUZ12-related accessions in pink and purple, respectively.

In the principal component analysis (PCA) of expression profiles, samples partitioned nicely with respect to disease and treatment condition, with the latter contributing the most to expression differences as reflected by PC1 (Supplementary Fig. 2). In line, the number of 492 differentially expressed genes (DEGs) along disease contrasts was about five times higher for contrasts assessing effects of hormonal treatments (Fig. 3a). The PCA further suggested great homogeneity in responses as the sample partitioning pattern with respect to disease status and person origin along PC2 remained highly similar for any of the three growth conditions reflected in PC1 (Supplementary Fig. 2). Moreover, cell type composition analyses of the samples against single cell reference data from uterus (Wang et al., 2020) and endometrial epithelial organoids (Fitzgerald et al., 2019) showed very consistent signatures with highest expression for epithelial cell types, as expected for the model and in line with the immunostainings for epithelial markers (Fig. 2; Supplementary Fig. 3 and 4).

### Transcriptional changes are partially shared between MRKH organoids and primary patient tissue

Focussing on the disease axis first, we identified 492 DEGs with 365 up- and 127 downregulated genes in expansion media (ExM) (Fig. 3b). Among these DEGs were twelve homeobox proteins (upregulated in MRKH: *LHX1, HOXD8, ONECUT3, LBX2, HOXB4, SATB1*, *HOXB6*; downregulated in MRKH: *EMX2, ZHX3, IRX3, NKX6-2, IRX5*). Of those, *LHX1*, *EMX2*, and the *Hox* genes have previously been associated with MRKH syndrome either by sequencing of patients’ blood or functional *in vivo* studies with animal models (Ledig et al., 2012, Masse et al., 2009, Miyamoto et al., 1997). Moreover, supporting the role of the 492 DEGs in Müllerian duct development, a large proportion of these genes are specifically activated during duct morphogenesis in chicken (Supplementary Fig. 5) (Roly et al., 2020).

Applying enrichment analyses to identify affected pathways and cellular processes as well as transcriptional regulators potentially driving the differential expression, we identified *plasma membrane* and *cell morphology* as most significant Gene Ontology (GO) terms and *ECM-receptor interaction* with respect to KEGG (Fig. 3c). A binding site analysis suggested highly significant overrepresentation of the *SP4* motif among DEG promotors (Fig. 3d). Interestingly, expression of *SP4* was decreased in MRKH organoids (Fig. 3e), agreeing with previous observations of a smaller uterus in *Sp4* knockout mice (Gollner et al., 2001). Complementary analyses that utilize ChIP-seq data and thereby account also for indirect binding events as well as transcription factors with less clear motifs (Puente-Santamaria et al., 2019) suggested the DEG set to be highly enriched for *EZH2* and *SUZ12* targets (Fig. 3f). Since *EZH2* has also been identified in our previous study of MRKH endometrial tissue (Hentrich et al., 2020), we see accumulating evidence that suggests further epigenetic investigations into the origins of the syndrome.

Similarity between MRKH organoids and the previously analysed endometrial tissue also existed with respect to DEGs. Comparing the set of 492 DEGs identified in MRKH organoids to the set of 2121 DEGs previously reported for MRKH endometrial tissue (Hentrich et al., 2020) led to 86 shared genes, of which 51 were up-respectively downregulated in both organoid and tissue (Fig. 4a). Among them were the *GATA5* transcription factor as well as the *HOXB4* and *HOXB6* homeodomain proteins, which were found up-regulated in MRKH patients in both studies (Fig. 4b).

**Figure 4.**
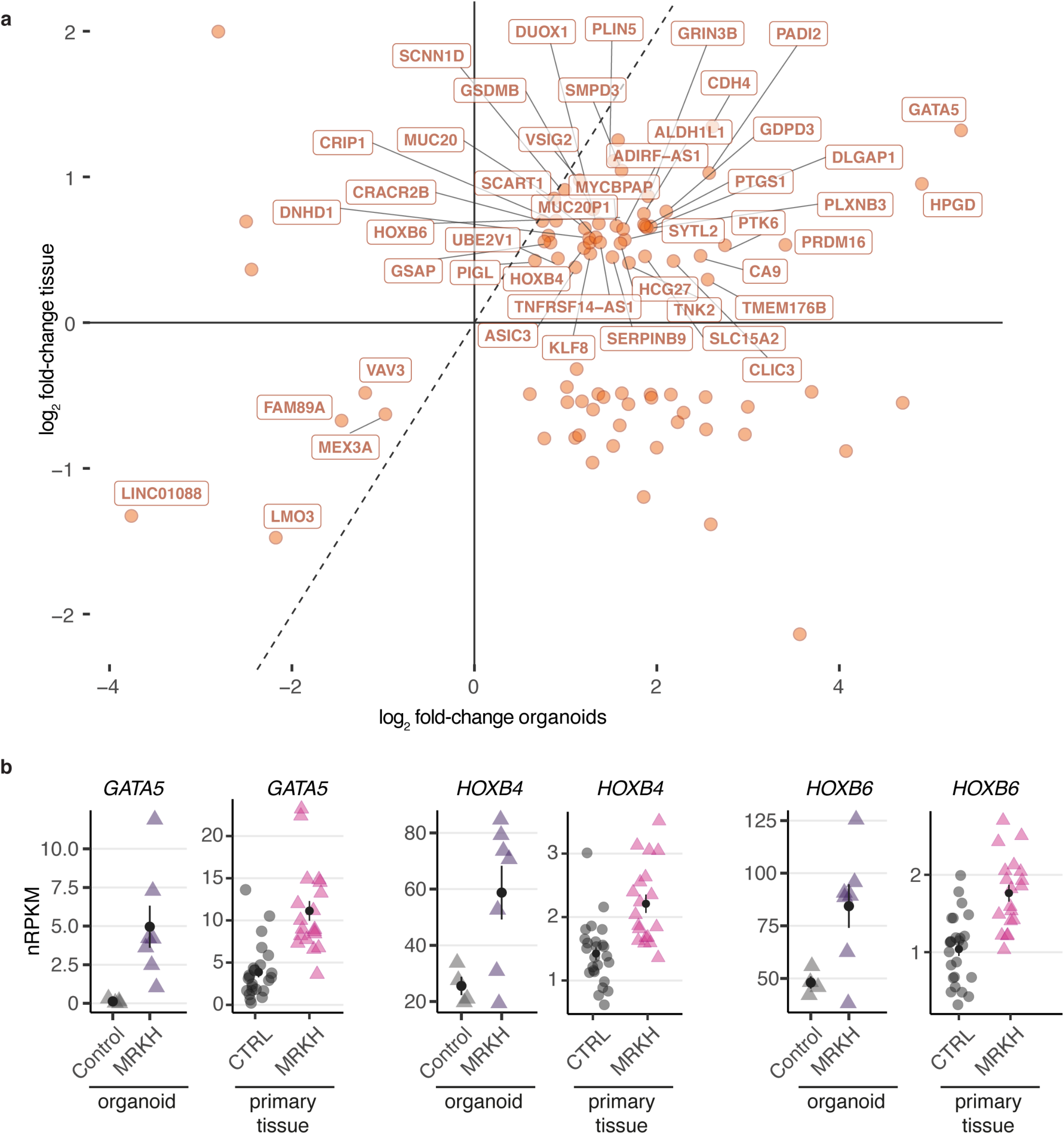
MRKH organoids share expression changes with endometrial tissue samples. **a)** Scatter plot of 86 common DEGs between MRKH organoids and endometrial patient tissue. DEGs with directional similarity (i.e. up-respectively downregulated in both tissue and organoid, 51 in total) are labelled. **b)** Expression levels of selected common DEGs plotted as individual data points with mean ± SEM in organoids as well as primary endometrial tissue (CTRL: unaffected women, MRKH: MRKH patients).

Despite the incomplete overlap of DEGs, which is likely attributable to the pure epithelial origin of organoids compared to the mixture of cell types (stromal, epithelial, endothelial and blood) found in primary tissue (Supplementary Fig. 3) (Hentrich et al., 2020), these results suggest the organoids to capture parts of the pathology in a highly homogeneous and reproducible way.

### Widespread transcriptomic response of endometrial organoids to hormonal treatments

Next, we sought to investigate transcriptional changes upon treatment with steroid hormones in endometrial organoids of MRKH patients and healthy controls as the human endometrium undergoes substantial remodelling mainly controlled by the steroid hormones E2 and P4 during menstrual cycle (Aghajanova et al., 2008, Roy and Matzuk, 2011). In order to investigate the hormone response of disease versus control organoids, RNA-sequencing data of E2- and P4-treated organoids were analysed for gene expression changes.

Under E2-treatment, 2321 DEGs were identified in control and 2290 DEGs in MRKH organoids (Fig. 5a), of which about three quarters overlapped (Fig. 5b). Furthermore, 594 of the DEGs have been described previously for endometrial epithelial organoids treated with E2 (Fitzgerald et al., 2019). As indicated by the PCA (Supplementary Fig. 2), the expression changes were highly homogenous between samples (Supplementary Fig. 6a) and encompassed genes such as *FOXJ1*, which activates genes essential for motile cilia formation and function, as well as *DYDC2*, a marker for ciliated cells (Supplementary Fig. 6b). Consistently, the most significant GO terms for these DEGs were *cilium organization* and *cilium assembly* (Fig. 5c). A fact that could also be observed by the immunostaining of hormone-treated organoids with the cilia marker acetylated alpha-tubulin (Fig. 5d). In addition, the expression of genes attributed to ciliated cells strongly increased upon treatment with E2 in both MRKH and control organoids (Supplementary Fig. 3 and 4).

**Figure 5.**
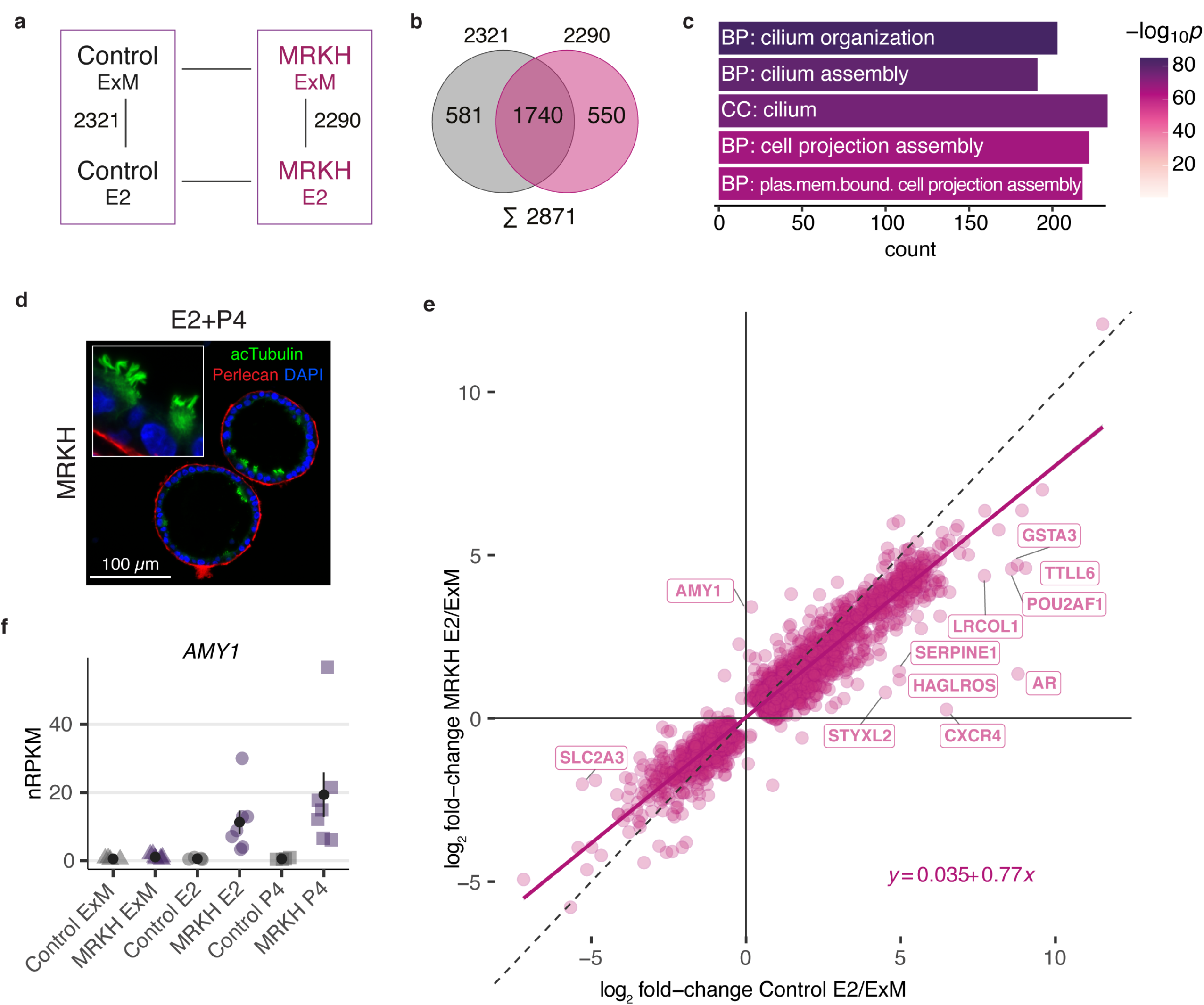
Widespread transcriptomic response of organoids upon hormonal treatment with beta-estradiol. **a)** Diagram of experimental groups kept in expansion medium (ExM) or treated with beta-estradiol (E2). Number of differentially expressed genes indicated for pairwise comparisons in control and MRKH organoids. **b)** Venn diagram comparing common and distinct DEGs upon beta-estradiol treatment between MRKH and control organoids. **c)** Enrichment analysis of overrepresented Gene Ontology terms among 2871 DEGs (union of DEGs comparing E2/ExM in MRKH and control organoids). Top five terms with number of associated genes shown according to their significance. CC: cellular compartment, BP: biological process. **d)** Representative immunofluorescence staining of a hormone stimulated MRKH organoid showing ciliated cells (green; acetylated α-tubulin). Perlecan-staining (red) was used to represent the epithelial polarity, nuclei shown with DAPI (blue), Scale bar = 100 μm **e)** Scatter plot of 2871 DEGs (union of DEGs contrasting E2/ExM) comparing expression changes in control (*x*-axis) versus MRKH (y-axis) organoids. DEGs that differ in their altered expression by more than | log_2_ FC | > 1 between control and MRKH are labelled. **f)** Expression levels of *AMY1* plotted as individual data points with mean ± SEM.

Intriguingly, despite largely similar gene expression changes upon E2 treatment between MRKH and control organoids, a few genes were specific to disease condition (Fig. 5e). Among them were genes such as *AMY1* with upregulation upon E2 treatment specifically in MRKH organoids (Fig. 5f). Consistently and previously unnoticed, *AMY1* also showed increased expression in MRKH endometrial tissue (Supplementary Fig. 6c). Hence, amylases – typically not expressed in endometrial tissue – might have been spuriously upregulated in MRKH patients, leading to tissue breakdown and degeneration.

In a next step, we studied the response of MRKH and control organoids to the combination of E2 and P4. As mentioned above, the DEG count was very similar and symmetric between E2 alone and the combination of E2 and P4 (Supplementary Fig. 7). In fact, for both control (Supplementary Fig. 7a) and MRKH organoids (Supplementary Fig. 7b), gene expression changes under the hormone treatments were almost identical as reflected by the nearly perfect standard diagonal (Supplementary Fig. 7c, d), and very few genes showed deviations (Supplementary Fig. 7e), indicating that P4 led to few additional transcriptome changes.

### Validation of disease- and patient-specific gene expression changes in endometrial organoids

Finally, we validated disease- and patient-specific gene expression changes of selected genes using quantitative PCR in the RNA-sequencing cohort as well as in an independent cohort (Fig. 6). We focused on the highly differentially expressed genes *LHX1* (associated in Müllerian agenesis (Huang et al., 2014)), *HOXD8* (homeobox gene highly expressed during the development of the chicken Müllerian duct(Roly et al., 2020)), *FAM3B* (a recently identified FGFR ligand implicated in posterior development (Zhang et al., 2021)), *NDN*, Androgen Receptor (*AR*) and *GATA5* (expressed during the Müllerian duct development and associated with abnormalities of the genitourinary tract of female mice (Roly et al., 2020, Molkentin et al., 2000)). Quantitative PCR assays nicely validated the RNA-sequencing results (Fig. 6b). Additionally, the expression changes found in our MRKH sequencing cohort were also found in the three MRKH models that were selected as an independent cohort (MRKH-IC) (Fig. 6b).

**Figure 6.**
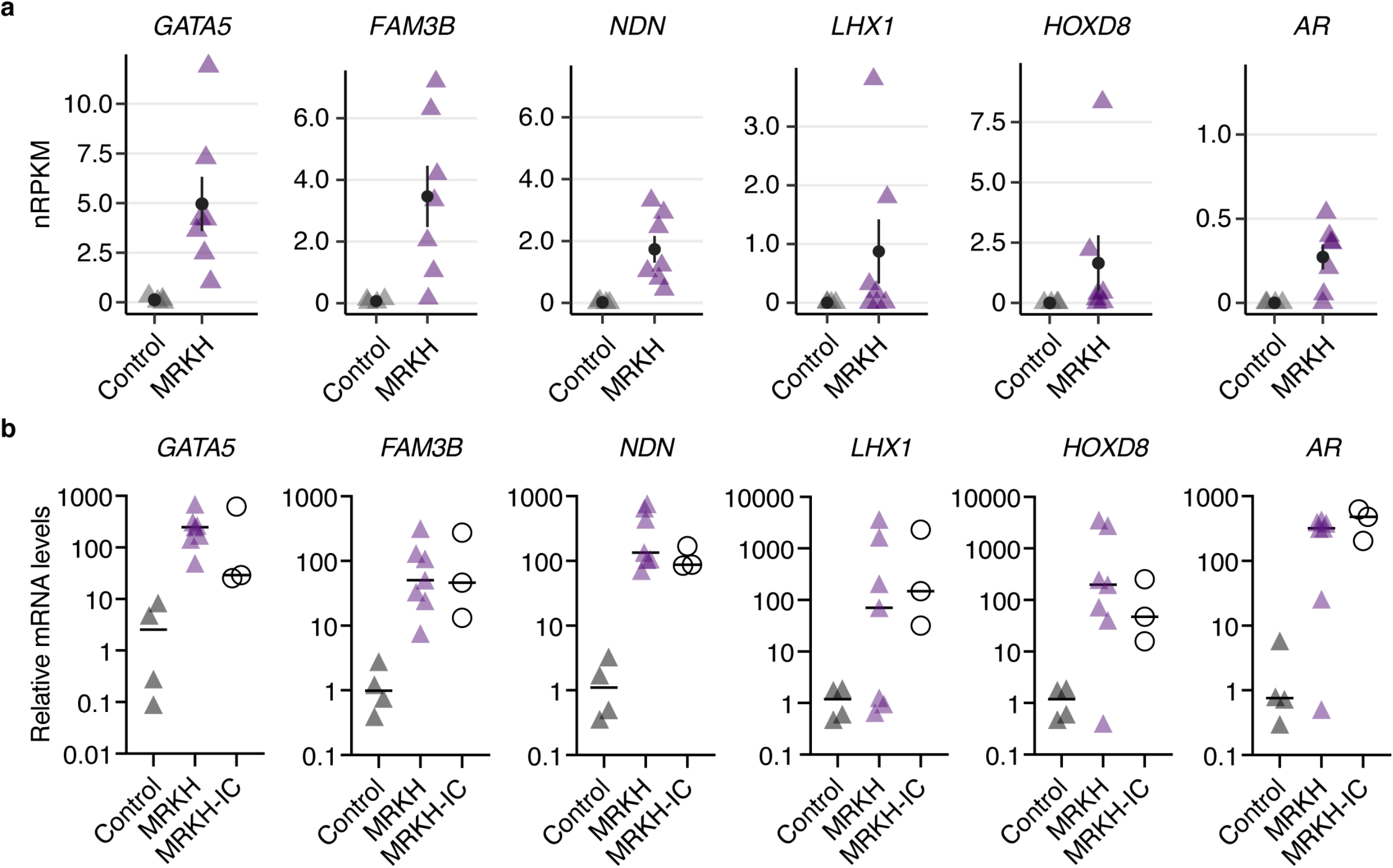
Validation of selected condition- and patient-specific gene expression changes. **a)** Expression levels of *GATA5, FAM3B, NDN, LHX1, HOXD8, and AR* plotted as individual data points with mean ± SEM based on RNA-sequencing data. **b)** Validation of expression changes seen in RNA-sequencing of the sequencing cohort (Control and MRKH) as well as an independent cohort (MRKH-IC) consisting of three models. Expression levels of *GATA5, FAM3B, NDN, LHX1, HOXD8, and AR* were investigated by qRT-PCR, using *SDHA* and *RPL13A* as reference genes. Data was normalized to control organoids to obtain relative mRNA levels and is shown as mean ± SEM. Each dot represents an individual organoid line.

## Discussion

During the last decade a plethora of research projects, mainly in the fields of genetics, have aimed at deciphering the cause of the multi-faceted MRKH syndrome (Herlin et al., 2020, Fontana et al., 2017). Reports of familial cases besides seemingly sporadic appearances have fueled hope to find a genetic cause of the disease. Yet, a common genetic determinant has remained elusive thus far. Sequencing studies are mainly performed on blood samples and the focus lays primarily on mutations in the patients’ germline. That, however, negates the possibility that the origin of MRKH syndrome lies in the developing tissue. Müllerian duct development (Santana Gonzalez et al., 2021) is, as all developmental processes, a highly complex sequence of events that involves activation and silencing of signaling pathways, expression of transcription factors (e.g., homeobox proteins), production of cascades of growth factors which all must be precisely coordinated in space and time to arrive at the desired outcome (Mammoto and Ingber, 2010). Disturbances in these pathways either by lacking factors or their deregulated expressions may, hence, lead to developmental defects, as seen in MRKH patients with uterus aplasia alone (MRKH Type I) or with associated malformations (MRKH Type II).

### MRKH organoids show no evidence of impaired hormone receptor function

To find potential disease-causing alterations, we strongly advocate for focusing on diseased tissue, rather than sequencing patient blood. So far, only one study reports the successful culturing of endometrial stromal cells isolated from MRKH rudimental uterine horns (Brucker et al., 2017). Here, we were able to isolate endometrium from rudimental uterine horns and establish long-term proliferating epithelial organoid cultures that share high similarity with healthy controls and are fully hormone responsive (Fig. 2 and 5). This is in stark contrast to the previously isolated endometrial stromal cells which showed impaired decidualization upon steroid hormone exposure (Brucker et al., 2017). We hypothesize that these results point to a pivotal interplay between stromal and epithelial cells might be of essence. *In vivo*, estrogen-stimulated stromal cells secrete growth factors (e.g., EGF and FGF-10) that lead to the proliferation of epithelial cells (Chung et al., 2015). Hence, impairment of estrogen receptivity in the stromal compartment might, in turn, lead to a reduction of these factors in MRKH patients and result in the reduced proliferation capacity seen *in vivo* but not in the organoid model due to the supplementation of these factors in the growth medium (Supplementary Table S2). Successful establishment and proliferation of endometrial cultures show that these cells are intrinsically able to proliferate once they are provided with external growth signals. These results suggest an essential crosstalk and interplay between stromal and epithelial cells in the pathogenesis of MRKH. The organoid model established here lends itself to further explore this and continue to explore the causal factors of the disease.

### Female reproductive tract development

During development of the female reproductive tract, a cascade of tightly regulated processes in the developing embryo need to take place. This is largely governed by tightly controlled transcription factor programs, including homeobox domain proteins. Clinically, MRKH represents a heterogeneous disease and is generally categorized into Type I (isolated finding) or Type II (accompanied by abnormalities of additional organ systems including mainly the kidneys and the skeleton) (Herlin et al., 2020). Interestingly, in our patient cohort about 90% of MRKH Type I cases have rudimentary horns, in contrast to Type II cases with only 38% (Supplementary Table S1). The multifaceted appearance of the uterine horns (absent, one vs. two, with endometrium and without, size variation) might point to defects at different phases of uterine development.

The uterus, both fallopian tubes, cervix and the upper third of the vagina develop from the Müllerian ducts (MD) whose development can be divided into three phases (Santana Gonzalez et al., 2021). The first phase is the specification phase when Müllerian precursor cells are specified in the surface epithelium at the cranial pole of the mesonephros (embryonic kidney). During phase 2, termed invagination, these cells start to become mesoepithelial and move between the mesenchyme in a caudal direction. In the final phase, the elongation phase, invaginating cells form the MD epithelium contact the Wolffian duct (WD) and the duct elongates. At this point, the most caudal part of the MD fuses into a single tube with a separating midline epithelial septum, forming the so-called midline uterovaginal canal between the two adjacent MDs which later disappears. The degree of midline fusion is vastly different between species. In mice, most of the MDs remain unfused and later give rise to large bilateral uterine horns instead of a single uterus in humans (Spencer et al., 2005). At this stage of development, the embryo is sexually indifferent and presents both with WDs and MDs (Kobayashi and Behringer, 2003). Under the influence of several signaling cascades and factors (e.g., Anti-Müller-Hormone; AMH) the MDs regress in males, whereas lack of these hormones in females leads to regression of the WDs (Kobayashi and Behringer, 2003). After female sex determination, expression of Hox genes and retinoic acid (RA) signaling drive segmentation of the proximal and distal Müllerian duct (Santana Gonzalez et al., 2021). Each specified Müllerian segment will give rise to a different part of the female reproductive tract with the most anterior differentiating into the oviducts and the most posterior into the upper vagina. We hypothesize, there is not a single defect in this finetuned developmental process that causes the MRKH syndrome but that the multifaceted clinical phenotype is owed to the fact that in a subset of patients one pathway was altered whereas in another subset a different one was. MRKH patients without rudiments and patients with two developed rudiments containing endometrium are very likely to have different alterations in their genome and/or epigenome. The latter seem to have had a caudal MD fusion defect whereas the former phenotype probably arises from a defect during elongation phase. The fact that MRKH patients have fallopian tubes (Herlin et al., 2020), both emerging from MDs, provides evidence that initial steps of MD development are still intact. Another possibility might be that regression of the WDs (driven by apoptosis and degradation of tissue (driven by metalloproteinases (Page-McCaw et al., 2007))) in MRKH patients was malfunctioned and led to partial regression of the adjacent MDs, which, in turn, lead to a fusion defect in the uterovaginal canal. Hence, the structure regressed and could not establish a contact with the urogenital sinus that is forming the lower part of the vagina. Investigation of endometrial tissue of patients with other Müllerian anomalies such as bicornuate uterus, uterus didelphys (defect in fusion) or a uterus septum (defect in septal resorption) might provide important insights in the causality of MRKH as these likely represent similar, but milder defects, compared to MRKH patients.

### Identification of disease-causing candidate genes in MRKH endometrial organoids

Uterine development occurs during the first trimester of development. At the time of surgery and tissue sampling, our patients have passed this point on average by 20 years (with one patient being already 41 years old) (Supplementary Table S1). It is remarkable that we were able to establish fully functional endometrial organoids from this tissue, and even more remarkable, that we were able to find specific expression differences in 492 DEGs including already suspected candidate genes such as *LHX1* and *HOX* genes. Surprisingly, these were mainly found expressed in MRKH organoids and absent in healthy controls. We speculate, this observation may explain why any single candidate gene may not necessarily be mutated in all patients. Mutations that cause a complete loss of function might have too severe developmental defects leading to embryonic lethality – and, hence, cannot be discovered. For example, homozygous *Lhx1* knock-out in mice leads to only a few neonates that are born with an MRKH-like phenotype, but also lack anterior head structures (Kobayashi et al., 2004). A deregulation (meaning not necessarily a knock-out) of this gene, specifically during uterine development or in specific regions of the developing female reproductive tract, might however cause MD anomalies while leaving other regions ‘unharmed’. The fact that a large portion of the MRKH candidate identification data is curated from knock-out studies in animal models (Masse et al., 2009), might be the reason why many of our identified genes have not yet been associated with the disease. The transcription factor GATA5 for example is highly expressed in MRKH endometrial organoids as well as in tissue (Hentrich et al., 2020) compared to healthy controls but has never been linked with MD anomalies although it is highly and dynamically expressed in the developing tissue as revealed by a large-scale RNA-sequencing study in chicken (Roly et al., 2020). In the future, shifting focus from knock-out studies towards interrogation of dysregulated gene expression such as CRISPR-A screens (Kampmann, 2018) might help to better understand the processes during uterine development. Several of our identified DEGs could serve as a starting point to select interesting candidates.

### The importance of identifying the cause of MRKH syndrome

The MRKH syndrome is the most common cause of uterine aplasia at a frequency estimated to be of 1 in 4,500 female newborns. Since the early 2000s, treatment of MRKH patients mainly focuses on the coexisting vaginal aplasia by creation of a neovagina to give patients the possibility of sexual intercourse (Rall et al., 2014). This vastly improves quality of life of affected women (Weijenborg et al., 2019). However, this treatment is not adequate to provide a solution for the missing reproductive ability in MRKH patients. Besides adoption and surrogacy, uterus transplantation has emerged as a highly promising method to overcome this shortcoming (Brannstrom et al., 2018). Both latter methods arrived with a plethora of ethical difficulties and researchers and clinicians are working urgently to find new ways of providing larger groups of patients with a cure. Organ-on-a-chip based methods as well as using animals or deceased patient uterus scaffolds represent opportunities for uteri reconstruction using patient-specific cells to overcome limitations of transplantation approaches (Bergmann et al., 2021). The development of these techniques in combination with an in-depth understanding of uterine development might mark a milestone for achieving a long-term treatment solution to some of the most severe uterine pathologies such as MRKH syndrome. To use patient-derived cells for uterus reconstruction, we need to get a deeper understanding of their functionality and their interactions. Hence, endometrial organoids from MRKH patients and their interaction with stromal cells need to be investigated further to achieve this goal. Our study shows that MRKH endometrial organoids are hormone responsive and show a high similarity to healthy endometrial epithelial cells. Future research in this emerging line of research, however, has to reveal whether this is sufficient for successful embryo implantation in a to-be-developed *de-novo* uterus from MRKH patient cells (Alzamil et al., 2021, Bergmann et al., 2021).

## Materials and Methods

### Patient cohort

All tissue biopsies were obtained from patients after informed written consent. The study was approved by the Ethics Committee of the Eberhard Karl University of Tübingen (Ethical approval 205/2014BO1 and 150/2018BO2) and is compliant with all relevant ethical regulations regarding research involving human participants. Rudimentary uterine horns were excised from 37 MRKH patients at the time of laparoscopically assisted creation of a neovagina and transported to the pathology lab in sterile containers. Uterine rudiments were sectioned perpendicular to the longest axis and macroscopically evaluated for presence of endometrial tissue. If present, endometrium was removed with as little attached myometrium as possible and further processed. For this study, endometrial tissue from twelve patients with MRKH syndrome was collected. Endometrial biopsy samples (obtained *via* Pipelle Endometrial Suction Curette (Medesign IC)) of four premenopausal patients served as controls. For full patient characteristics see Supplementary Table 1.

### Processing of endometrium samples and organoid culture setup

Tissue samples were minced into small pieces (1-3 mm^3^) using a scalpel and dissociated with collagenase/dispase (1 mg mL^-1^; COLLDISP-RO, Roche) in the presence of Rock inhibitor (RI; Y-27632, 10 μM; M1817, Abmole Bioscience) for 1 h at 37°C on a shaking table. The digestion was attenuated by addition of medium (advanced DMEM/F12 - without serum) and centrifuged at 478 g for 10 min. The final pellet was resuspended in advanced DMEM/F12 supplemented with 1% Glutamax/1% HEPES/1% penicillin/streptomycin (all Gibco) and the desired amount of cell suspension mixed with Basement Membrane Extract (Type 2, 3533-001-2, Trevigen; BME) at a ratio of 65% BME to 35% cell suspension. Twenty μL droplets were plated out on pre-warmed 48 well plates and placed upside-down in an incubator (37°C, 5% CO_2_) for solidification. Afterwards culture medium (Supplementary Table 2) was added to each well and renewed every 3 days. Noggin conditioned medium from HEK293T-Noggin-Fc-cells (kindly provided by Hans Clevers, Utrecht, Netherlands) was produced as previously described(Farin et al., 2012). R-Spondin conditioned medium was produced with Cultrex^®^ HA-R-Spondin1-Fc 293T Cells (Trevigen) according to the distributors protocol.

### Passaging/cryopreserving of organoid cultures

For passaging or cryopreservation, organoids were recovered by resuspending the BME-drops in ice-cold advanced DMEM/F12 and transferred to 15 mL tubes. The organoid suspension was either mechanically or enzymatically (25% 1xTrypLE Express (Gibco)/75% advDMEM/F12) dispersed and then pelleted. For further culture the pellet was reconstituted in advanced DMEM/F12 and mixed with BME at a ratio of 65% BME to 35% cell suspension and cultured as described above. For cryopreservation, the cell pellet was resuspended in Recovery™ Cell Culture Freezing Medium (Gibco), the solution transferred to cryo vials and then cooled down in CoolCell™ LX Freezing Containers (Merck) in a −80°C freezer. The next day, vials were transferred for long-term storage to liquid nitrogen tanks.

### Paraffin sections and immunohistochemistry

Tissue and organoids were fixed in 4% paraformaldehyde followed by dehydration, paraffin embedding, sectioning (4 μm), and standard H&E staining. Immunohistochemistry was performed on a Ventana Discovery automated immunostaining system (Ventana Medical Systems, Tucson, USA) using antibodies as specified in Supplementary Table 2.

### Immunofluorescence

For characterization of organoids, FFPE-sections (4 μm) were subjected to heat induced antigen-retrieval and incubated with primary antibodies as specified in Supplementary Table 2. Sections were incubated overnight in a humidified chamber at 4°C and afterwards washed three times with PBS containing 0.1% Tween-20 and incubated for 1 h at room temperature with the respective secondary antibody (Supplementary Table 2). Finally, sections were again washed and mounted in ProLong Diamond Antifade mounting media containing DAPI (Thermo Fisher Scientific). Images were acquired with the EVOS M7000 imaging system and processed using Image J (NIH, USA).

### Hormone treatment of organoid cultures

Organoids were passaged as described above and allowed to grow for 4 days in standard culture medium (<u>E</u>xpansion <u>M</u>edium; ExM). Afterwards three different groups were established: Group 1 (ExM) served as an untreated control sample and received ExM for an additional 6 days, with the medium refreshed every other day. Group 2 (E2) was cultured with ExM containing 10 nM beta-estradiol (Sigma, Cat. No. E8875) for 6 days. Again, the medium was renewed every other day. Group 3 (P4) first received ExM supplemented with 10 nM beta-estradiol for 2 days and afterwards ExM with 1 μM progesterone (Sigma, Cat. No. P8783) and 1 μM cAMP (Tocris, Cat. No. 1140) in addition to 10 nM beta-estradiol for additional 4 days. After a total of 10 days, all organoids were harvested. For this, medium was removed from the wells and the BME-domes incubated with 1xTrypLE Express at 37°C for 10 min. PBS was added to dilute the TrypLE Express and the suspension centrifuged at 478 g for 10 min. After discarding the supernatant, the pellet was resuspended in PBS and centrifuged again to remove leftover BME. After removing the supernatant, cell pellets were snap frozen in liquid nitrogen and then stored at −80°C.

### RNA isolation and sequencing

Total RNA was isolated from organoid cultures using the RNeasy Mini Kit (74004, Qiagen). Simultaneous elimination of genomic DNA was achieved with on-column DNA digestion (79254, Qiagen) according to the manufacturer’s protocol. Quality was assessed with an Agilent 2100 Bioanalyzer. Using the NEBNext Ultra II Directional RNA Library Prep Kit for Illumina and 100 ng of total RNA for each sequencing library, poly(A) selected paired-end sequencing libraries (101 bp read length) were generated according to the manufacturer’s instructions. All libraries were sequenced on an Illumina NovaSeq 6000 platform at a depth of around 40 mio reads each. Library preparation and sequencing procedures were performed by the same individual, and a design aimed to minimize technical batch effects was chosen.

### Quality control, alignment, and differential expression analysis

Read quality of RNA-seq data in fastq files was assessed using *FastQC* (v0.11.9) (Andrews, 2010) to identify sequencing cycles with low average quality, adaptor contamination, or repetitive sequences from PCR amplification. Reads were aligned using *STAR* (v2.7.6a) (Dobin et al., 2013) allowing gapped alignments to account for splicing against the *H. sapiens* genome from *Gencode* v35. Alignment quality was analyzed using *samtools* (v1.10) (Li et al., 2009). Normalized read counts for all genes were obtained using *DESeq2* (v1.32.0) (Love et al., 2014). Transcripts covered with less than 50 reads (median of all samples) were excluded from the analysis leaving 15,413 genes for determining differential expression. Cut-offs of |log_2_ fold-change| ≥ 0.5 and BH-adjusted *p*-value ≤ 0.05 were set to determine differentially expressed genes. Gene-level abundances were derived from *DESeq2* as normalized read counts and used for calculating the log_2_-transformed expression changes underlying the expression heatmaps for which ratios were computed against mean expression in control samples. The *sizeFactor*-normalized counts provided by *DESeq2* also went into calculating nRPKMs (normalized Reads Per Kilobase per Million total reads) as a measure of relative gene expression (Srinivasan et al., 2016).

### Gene annotation, enrichments, and regulator analyses

*G:Profiler2* (v0.2.0) was employed to identify overrepresented Gene Ontology terms for differentially expressed genes (Raudvere et al., 2019). Transcription factor binding site analyses were carried out in *PScan* (v1.6) (Zambelli et al., 2009) on the *H. sapiens* genome considering −450 to +50 bp of promoter region for motifs against the JASPAR 2020_NR database. *TFEA.chip* (v1.12.0) was employed with default parameters to determine transcription factor enrichments using the initial database version of ChIP-Seq experiments (Puente-Santamaria et al., 2019). Cell type-specific endometrial marker genes were taken from a study with single cell profiles from uterine tissue (Wang et al., 2020) as well as endometrial epithelial organoids (Fitzgerald et al., 2019).

### Reverse transcription quantitative PCR (RT-qPCR)

Total RNA was extracted from organoids using RNeasy Mini Kit (74004, Qiagen), simultaneously eliminating genomic DNA with on-column DNA digestion (79254, Qiagen). Equal amounts of total RNA (1 μg) were reverse transcribed using the Maxima™ H Minus cDNA Synthesis Master Mix (M1661, Thermo Fisher Scientific). To analyze gene expression, 5 ng cDNA was subjected to real time qPCR using PowerUp™ SYBR^®^ Green Mastermix (A25741, Thermo Fisher Scientific) on the QuantStudio 3 Real-Time PCR system (A28567, Thermo Fisher Scientific). Thermal cycling was performed with 3 min at 95°C, followed by 40 cycles of 95°C for 15 s and 60°C for 60 s. The specificity of the RT-qPCR products was assessed by melting curve analysis. Relative quantification was performed using the 2^−ΔΔCt^ method with *SDHA* and *RPL13A* as reference genes. Expression was normalized to the endometrial control group. All experiments were performed in duplicates. PCR primers are listed in Supplementary Table 4.

## Acknowledgements

We thank all patients who participated in the study. Special thanks go to Ingrid Teufel and Sabine Hofmeister for excellent technical support. We thank the team of pathologists of the Department of Anatomic Pathology, University of Tübingen, and the NGS Competence Center Tübingen, foremost Nicolas Casadei and Olaf Rieß, for their support. We thank Nico Weber for his input in the initial analysis of the RNA-sequencing data. Special thanks goes to Ana Maia dos Santos Leite for her critical input throughout the study and for her valuable comments on the manuscript.

## Author contribution

A.K. conceived and designed the project. A.K., A.R.S., K.R., S.Y.B. supervised the experimental work. T.H. and J.S-H. performed the bioinformatic analysis of RNA-sequencing data. K.R. and S.Y.B. obtained patient samples. M.P. and A.K. processed patient samples. N.W. performed the RT-qPCR validation. A.K., M.P., T.H. and J.M.S.H. wrote the manuscript with input from all authors. All authors read and approved the final manuscript. Part of the presented work here is part of the doctoral thesis of M.P.

## Data availability

RNA-sequencing data that support the findings of this study have been deposited in the European Genome-phenome Archive (EGA) under primary accession [pending submission validation].

---

***Supplementary Figure 1. Long-term 3D organoid cultures can be established from healthy endometrium***

**a)** Endometrium biopsy of a healthy control patient obtained with a Pipelle. Equal parts were used for tissue digestion and subsequent organoid setup as well as pathological characterization; Scale bar = 10 mm

**b)** Immunohistochemical characterization of control endometrium. Analyzed was Estrogen receptor alpha (ER), Progesterone receptor (PR), and proliferation marker Ki67/Mib1. The tissue shows endometrial gland structures with widespread and intense ER and PR expression in glandular and stromal compartments. Proliferative capacity (Ki67/Mib1) can be seen by in stromal and epithelial cells. Scale bar = 500 μm

***Supplementary Figure 2. Organoid expression profiles reflect partitioning of samples according to condition and treatment*.**

Principal component analysis of gene expression profiles for all samples based on the top 500 most variable genes. Axis percentages indicate variance contribution of the first two principal components. Control: organoids derived from unaffected women, MRKH: organoids derived from MRKH patients; ExM: organoids grown in expansion medium, E2: organoids treated with beta-estradiol, P4: organoids treated with beta-estradiol and progesterone.

***Supplementary Figure 3. Epithelial cells dominate cell type-specific expression signature in organoids*.**

Cell type-specific gene expression per sample for unciliated and ciliated epithelium, endothelium, stromal fibroblasts, lymphocytes, and macrophages. Boxplots show geometric mean as well as 10th, 25th, 75th, and 90th quantile of expression values for all genes grouped based on single-cell reference data of human endometrium(Wang et al., 2020).

***Supplementary Figure 4. Ciliated cell-specific genes increase expression upon treatment with steroid hormones*.**

Cell type-specific gene expression per sample for unciliated, ciliated, epithelium, proliferative, and stem cells. Boxplots show geometric mean as well as 10th, 25th, 75th, and 90th quantile of expression values for all genes grouped based on single-cell reference data of endometrial epithelial organoids of control samples(Fitzgerald et al., 2019).

***Supplementary Figure 5. Several differentially expressed genes play a role in the embryonic chicken Müllerian duct*.**

Z-score heatmap of 85 differentially expressed genes also altered in Müllerian duct development in chicken. Of 492 DEGs, 251 orthologues genes were found in chicken and of those 85 were differentially expressed and had a CPM >5 in one of the static or dynamic transcriptomic changes during duct formation pointing at developmentally regulated genes(Roly et al., 2020). Left part of the heatmap reflects transcriptomic changes in chicken based on data from Roly *et al*. and right part shows expression changes of these genes observed in MRKH versus control organoids.

***Supplementary Figure 6. Similar transcriptomic response to beta-estradiol in MRKH and control organoids*.**

**a)** Expression profiles (log_2_ expression change relative to control ExM group) of 2871 DEGs (union of DEGs comparing E2/ExM in MRKH and control organoids) across all samples. Rows hierarchically clustered by Euclidian distance and *ward.D2* method. Patient origin color-coded.

**b)** Expression levels of four genes showing pronounced up- (*FOXJ1, DYDC2*) respectively downregulation (*KRT4, VGLL1*) upon treatment with beta-estradiol. Plotted as individual data points with mean ± SEM.

**c)** Expression levels of *AMY1* in primary endometrial tissue plotted as individual data points with mean ± SEM (Hentrich et al., 2020).

***Supplementary Figure 7. Marginal differences of transcriptomic response to combined beta-estradiol/progesterone treatment versus beta-estradiol alone*.**

**a)** Number of differentially expressed genes between control groups upon treatment with beta-estradiol (E2) or progesterone (P4). Venn diagram comparing common and distinct DEGs of pairwise comparisons in the right panel.

**b)** Number of differentially expressed genes between MRKH groups upon treatment with beta-estradiol (E2) or progesterone (P4). Venn diagram comparing common and distinct DEGs of pairwise comparisons in right panel.

**c)** Scatter plot of 2529 DEGs (union of DEGs in A) depicting expression changes of E2/ExM (*x*-axis) and (P4/ExM) (y-axis) in control organoids. DEGs differing between both treatments by more than |log_2_ FC | > 1 are labelled.

**d)** Scatter plot of 2657 DEGs (union of DEGs in B) depicting expression changes of E2/ExM (*x*-axis) and (P4/ExM) (y-axis) in MRKH organoids. DEGs differing between both treatments by more than |log_2_ FC| > 1 are labelled.

**e)** Expression levels of selected genes showing pronounced differential expression upon treatment with beta-estradiol (E2) and progesterone (P4), respectably. Plotted as individual data points with mean ± SEM.

***Supplementary Table 1. Demographic and clinical characteristics of MRKH patients*.**

***Supplementary Table 2. Culture medium for expansion of control and MRKH organoids*.**

***Supplementary Table 3. List of antibodies used for immunostaining*.**

***Supplementary Table 4. List of primers for qPCR*.**

***Supplementary Table 5. Differentially expressed genes for all primary contrasts*.**

